# Metabolic engineering of *Bacillus subtilis* with an Endopolygalacturonase gene Isolated from *Pectobacterium. carotovorum*; a Plant Pathogenic Bacterial Strain

**DOI:** 10.1101/2021.08.17.456673

**Authors:** Nagina Rafique, Saiqa Bashir, Muhammad Zubair Khan, Imran Hayat, Willium-Orts, Dominic W. S. Wong

## Abstract

Pectinolytic enzymes (pectinases) produced by microbes are highly important for their biotechnological use in processing of vegetables and fruits beverages and use in pulp and paper industry. A pectinase, namely endo-polygalacturonase (endo-PGase), encoding gene isolated from *Pectobacterium carotovorum*, a plant pathogenic strain of bacteria was successfully cloned into a secretion vector pHT43 having σ^A^-dependent promoter P *grac*. For enhanced expression analysis, competent cells of *Bacillus subtilis* (WB800N) were prepared at stationary phase using high salt medium. The recombinant *B. subtilis* competent cells, harbouring the engineered pHT43 with the endo-PGase gene were cultured in 2X-yeast extract tryptone medium. The recombinant endo-PGase enzyme was secreted directly into the medium after 72 hours of the first IPTG induction. The recombinant endo-PGase was screened for its activity at various temperatures and pH ranges. Optimal activity was found at pH 5.0 and a temperature of 40°C with a stability ranging from pH 5.0-9.0. For detection of metal ion effect, recombinant enzyme was incubated with 1mM concentration of; Ca^++^, Mg^++^, Zn^++^, EDTA, K^++^ for 45 minutes. Resultantly, Ca^++^, EDTA and Zn^++^ strongly inhibited the enzyme activity. The chromatographic analysis of enzymatic hydrolysate of polygalacturonic acid (PGA) and pectin substrates using HPLC and TLC revealed that tri and tetra-galacturonates were the end products of hydrolysis. The study led to the conclusion that endo-PGase gene from the plant pathogenic strain was successfully expressed in *Bacillus subtilis* and could be assessed for enzyme production using a very simple medium with IPTG induction. These findings proposed that the *Bacillus* expression system might be safe for commercial enzyme production as compared to yeast and fungi to escape endotoxins.

## Introduction

In microbial world, the genus *Erwinia* consists of plant pathogenic bacterial species that cause diseases in numerous plants. These bacteria cause disease in plants by producing high quantities of several types of cell-wall-degrading-enzymes collectively called exoenzymes such as; pectinases, proteases, cellulases etc which catalyze the cell-wall-breakdown leading to release of plant nutrients for the growth of these bacteria [1]. Among these, *Erwinia carotovora* synonymously also known as *Pectobacterium carotovorum* [2] a gram-negative species, belonging to family *Pectobacteriaceae*, is a plant pathogen to a wide range of agriculturally and economically important plants. It produces pectinolytic enzymes that hydrolyse pectin-polysaccharides within plant cells. It is a very economically important plant pathogen in terms of postharvest losses and causes decay in stored fruits and vegetables [3]. The virulence factors of *Pectobacterium* are grouped under pectinases which include; pectate lyase (*Pel*), pectin lyases (*Pnl*), and pectate hydrolases (*Peh*) also called polyglacturonase. Among these pectinases, pectate hydrolase (*Peh*) which is also called polyglacturonase cause widespread maceration of tissue, rotting, and consequently death of the entire plant [3, 4, 5].

Pectinases have numerous industrial applications associated with processing of natural products for extraction purposes [6]. Pectinases are involved in depolymerisation of pectin by its hydrolysis, trans-elimination and de-estrification reactions. Endo-polygalacturonase (EC 3.2.1.15), exo-polygalacturonase (EC 3.2.1.67), pectate lyase, pectin lyase and pectin esterase are among these pectinases reported in literature [6,7].Among these pectinases, endo-polygalacturonase (EndopGase) has been extensively studied for random hydrolysis of α-1, 4 glycosidic bond in the linear chain of pectin [8, 9] Pectin is degraded by cleavage within α-galacturonic acid polymer structure by the action of endo-polygalacturonases (EC 3.2.1.15) [10].Because of wide applications in food, feed, paper, fruit juice and textile industries, endo-polygalacturonase has gained significant research attention in recent years [6]. Studies on pectinase of microbial bioresources reported 25% of its share for global food and that the industrial enzymes marketing and sales are increasing constantly [11]. Moreover, there is a global market projection of enzymes reaching 6.3 billion USD in the current year 2021 [12]. In food industries, pectinases such as polyglacturonases play a pivotal role for that these are used in extraction of fruit-juices, clarification of wines, cocoa, tea, concentration of coffee; fermentation; extraction of vegetable –oil, pickling and processing of jams and jellies [13, 14]. Additionally, pectinases are used in pulp, paper and fiber industries, treatment of waste-water, poultry-feed additives and biofuels productions [14, 15, 16]. Enzymatic catalysis of bioresources is affected by many other factors such as; pH, temperature, nitrogen and carbon source, incubation time, agitation, substrate type and its concentration and utilization of various enzyme formulations during biotechnological processing [15, 17, 18].

Meachanism of Catalytic reaction is shown in the following equation:

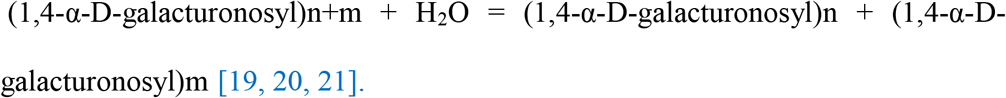

Many endo-PGs are recombinantly expressed in *E. coli* [22] and also in *Pichia pastoris* [23, 24]. Bacterial strains other than *E. coli* are becoming more applicable for heterologous protein expression. Gram positive bacillus strains are comparatively remarkable alternative for gene expression than that of *E. coli. Bacilli* are more beneficial as they have no lipopolysaccharides in the outer layer of cell wall and thus there is no danger of endotoxins production. Because of high secretion capacity and direct export of protein into medium *Bacillus* strains have become the most stimulating host system. Among Bacilli, *B. subtilis* is a well-studied prokaryotic strain and considerably used for protein expression [25]. Previously we cloned and expressed endo-PGase gene from *P. carotovorum* into *Pichia pastoris* [26]. with encouraging results. Here for the first time we report the expression of an endo-PG gene from *Pectobacterium carotovorum* into *Bacillus subtilis*. The recombinant enzyme has been characterized by performing different molecular and biochemical analyses.

## Materials and Methods

### Bacterial strains, plasmids, bacterial culture media and Reagents

The bacterial strains *E. coli* JM109 were purchased from Invitrogen life technologies for initial cloning and of *peh* gene. For gene expression, *Bacillus subtilis* (WB800N) and its corresponding pHT43 vector (Fig 1) for enzyme production and secretion [27] were purchased from MoBiTech (Molecular Biotechnology, GmbH, Germany). Qiagen miniprep Kit`for recombinant plasmid isolation and purification and SDS-NuPAGE precast gels for molecular characterization of expressed enzyme were obtained from Invitrogen (Carlsbad, CA). Polygalacturonic acid (PGA) sodium salt and citrus peel pectin were purchased from Sigma-Aldrich (MO, USA). Chemicals for culture medium and agar were gathered from Difco Laboratories (Detroit, MI). All other chemical/reagents used in molecular characterisation, TLC, HPLC and enzyme activity bioassays were of analytical grade.

**Fig 1.** pHT43 an IPTG-inducible expression vector for *Bacillus subtilis*. pHT43 was manipulated for the rapid secretion and purification of recombinant proteins in *Bacillus subtilis*. pHT43and the eight-fold protease-deficient *B. subtilis* strain WB800N, for secretory protein production are available from MoBiTec GmbH (Göttingen) [27].

### Gene Synthesis, Plasmid construction and transformation of *Bacillus subtilis*

Endo-polygalacturonase *peh* gene was synthesized by Genscript (Piscataway, NJ) using sequence data of the *peh* gene of *Pectobacterium carotovorum* from NCBI and DDBJ. A fragment of 1209bp open reading frame was amplified by deducing the desirable gene sequence using accession numbers; X52944 genomic DNA translation: CAA37119.1, and L32172 genomic DNA translation: AAA57139.1 The amplified gene fragment was first ligated in the modified pUC57 vector. Then the gene was subcloned in fusion to the amyQ signal peptide downstream of the p_*grac*_ promoter in *Bacillus* pHT43 vector (MoBiTec GmbH, Germany) using codon optimisation approach.

Briefly, the endo-PGase gene was PCR amplified and a secretion vector pHT43 specific to *Bacillus subtilis strain WB800N* was constructed based on σ^A^-dependent promoter P*grac*. The constructed vector consisted of *gro*E promoter, SD_*gsiB*_ gene sequence and a SP_*amyQ*_ (*Bacillus amyloliquefaciens* signal peptide) with multiple cloning site (*Bam*HI, *Xba*I, *Aat*II and *Sma*I) and *lac*O operator. The expression vector was transformed into *E*.*coli* JM109 before transforming into *Bacillus subtilis* following instructions from manual provided by MoBiTec GmbH, Germany. The transformed colonies were screened by double digestion of isolated plasmids from overnight grown cultures and positive transformants were confirmed by agarose gel electrophoresis and DNA sequencing simultaneously.

### Genetic techniques: Preparation of Competent cells

*Bacillus subtilis* competent cells were prepared by inoculating 50 ml HS medium with overnight grown culture of appropriate recipient cell line of *Bacillus subtilis* and incubated at 37 °C shaker incubator. The growth was recorded after every 40 min and at stationary phase 10 ml of sample was taken with the interval of 15 min, sterilized glycerol stock (87%) was added and placed on ice for 15 min. The entire sample was fractionated into aliquots of 1 ml and was frozen using liquid nitrogen following the instruction manual supplied by MoBiTec GmbH, Germany (available online). The prepared stocks of competent cells (*Bacillus subtilis* WB800N) were stored at -80 °C freezer. Desirable stocks quantities were shifted to -30°C freezer and gradually thawed in ice for transformation by recombinants as and when scheduled.

### Recombinant DNA techniques: Expression of Recombinant Endo-polygalacturonase

The enzymatic protein was expressed by inoculating fresh liquid culture at OD 0.8 with 1mM IPTG following the second induction of IPTG which was done after 8 h of the first induction. The culture was grown at 37 °C in a shaking incubator at 225 rpm. Endo-polygalacturonase production from recombinant *Bacillus subtilis* was first tested in 250 ml flask with 50 ml of 2X-YT medium. Cell free culture supernatants were harvested by centrifugation after 24h of the first induction and up to 120 h, the samples were continuously collected after every 24 h and assayed for endo-PGase activity following DNS method [28].

### Biochemical Characterization of Recombinant Endo-PGase

In order to determine the optimal pH and temperature requirements of recombinant endo-PGase, the enzyme activity was measured at pH ranging from 4.0-10.0 and temperature ranging from 20 °C - 70 °C. For estimation of pH stability, enzyme samples were incubated with buffers of different pH at 25 °C for 4 h and then the residual activity was measured. Thermal stability was evaluated by incubating the enzyme samples at temperatures ranging from 20 °C to 70 °C in 0.2M acetate buffer of pH 5.0 for 45 min and residual activity was measured by DNS method following Wong et al [28]. Effect of five different metal cations was determined by incubating enzyme samples with 1 mM concentration of each metal ion i.e. Ca^++^, Mg^++^, EDTA, Zn^++^ and K^++^. At the same time one control sample without having any metal cations was also included. The enzyme activity was measured by DNS method as previously described [26, 28].

### Molecular Analysis of Recombinant Endo-PGase Gene Expression

Cell free culture supernatant was analysed for recombinant protein expression with SDS-PAGE electrophoresis system. SDS-PAGE gel electrophoresis of cell free supernatant was carried out using Novex mini gel system (Invitrogen) with Nu-PAGE 10 % Bis-Tris gels and MES-SDS running buffer. A known molecular weight (kDa) protein standard marker was loaded in the first well to quantify the unknown weight of the recombinant enzyme samples loaded in the adjacent wells. Gel was stained with Coomassie Brilliant Blue, destained and then photographed.

### Enzyme activity analysis

Culture supernatant was tested for enzyme activity using solid as well as liquid assay methods. For plate assay the aliquots of enzyme were inoculated into wells on plates containing 0.5 % pectin (from citrus peel) and 0.5 % agarose. Plates were incubated overnight at 37 °C and stained with 0.02 % ruthenium red. Liquid assay of culture supernatant was performed using DNS method to measure the galacturonic acid produced [28] The reaction mix contained 75 µl of polygalacturonic acid (PGA) sodium salt (1%), 75 µl of 0.2M sodium acetate buffer of pH 5.0 and different concentrations of enzyme about 150 µl. The reaction mix (pH 5.0) was incubated at 40 °C for 1 h and unit of enzyme activity was calculated as µg of galacturonic acid/min at 37 °C temperature.

### TLC and HPLC Analysis of Hydrolysis products

Analysis of hydrolysis reaction products of PGA-sodium salt was performed using thin layer chromatography. An aliquot of 10 µl reaction was spotted onto TLC plates (20 × 10 cm silica plate) with mobile phase containing 2:1:1 ratio of ethyl acetate/acetic acid/water. The hydrolysis reaction products and standard oligo-galacturonates of different pectic substrates were analysed by HPLC using Zorbax-SAX column (200 ×10.0 mm, Agilent) in 0.3M sodium acetate (pH 5.0) at a flow rate of 0.9 ml per min at 40 °C [29]. The injection volume of 5 µl was applied to the injector and monitored by detector. Organic acids peaks were screened using refractive index detector.

#### Bioinformatics and Molecular Graphics

For recombinant vector construction and sequence analysis, Geneious was used [30]. Multiple sequence alignment was performed using by ClustalW and graphically presented by Bioedit. A phylogenetic tree was constructed with the help of Jalview using Neighbor-Joining for the generation of dendrogram.

For recombinant vector construction and sequence analysis, Geneious was used [30]. Multiple sequence alignment was performed using by ClustalW and graphicalyl presented by BioEdit Sequence Alignment Editor [31]. Evolutionary analyses were conducted in MEGA (Molecular Evolutionary Genetics Analysis) 7 [32]. The evolutionary history was inferred using the Neighbor-Joining method [33]. The evolutionary distances were computed using the Poisson correction method [34] and are in the units of the number of amino acid substitutions per site. The analysis involved 9 amino acid sequences. All positions containing gaps and missing data were eliminated. There were a total of 342 positions in the final dataset. 3D structure of EndoPGase enzyme was visualized with the Phyre2 web portal for protein modeling, prediction and analysis [35].

### Statistical analysis

For plotting graphical figures and calculations of standard bars Kaleidah Graph software was used [36].

## Results and Discussion

### Cloning, Isolation and Characterisation of *PehA* gene

In this study an endo-peh encoding gene of *Pectobacterium carotovorum* origin was successfully cloned, characterised, ligated into the multiple cloning site of vector pHT43-amyQ (Fig 1) and expressed in *Bacillus subtilis* (WB800N).

The desired PCR product was 1209bp first recovered by agarose gel electrophoresis (Fig 2). The isolated gene was further confirmed by sequence analysis.

**Fig 2.** Horizontal gel electrophorogresis of the cloned *PehA* gene from *P. carotovorum*. Double digestion with BamHI and XbaI for the clone confirmation of transformed colonies (left). Restriction digestion with Cla. Lane 1, 7 and 9 represents the clone confirmation of our gene of interest (right).

The open reading frame of cloned gene confirmed the encoded polypeptide of 402 amino acids. The signal peptide, as confirmed from Phyre2 showed signal peptide of 26 amino acid residues with the N-terminus. BLAST, PDB and Uniprot search for the amino acid sequence revealed close evolutionary relation of it with the isolated gene. Multiple sequence alignment of related EndoPGase across selected microbial and Plant genera with the isolated gene is shown in (Fig 3) which showed sequence identity of 97.51% with *Pectobacterium brasiliense* (WP_039510994.1), 62.50% with *Erwinia tasmaniensis* (WP_042958805.1), 57.65% with *Zymobacter palmae* (WP_027705705.1), 50.00% with *Pantoea ananatis* (PA13 AER33559.1), 32.54% with *Solanum lycopersicum* (AAB09576.1), 32.37% with *Zea mays*, (ACF85710.1), 31.50% with *Prunus persica* (AEI70578.1) and 28.23% with *Aspergillus niger* (CAA74744.1) respectively.

**Fig 3.** Multiple sequence alignment of *PehA* gene with the selected taxa. (from top to bottom); *Pectobacterium carotovorum, Pectobacterium brasiliense* (WP_039510994.1), *Erwinia tasmaniensis* (WP_042958805.1), *Zymobacter palmae* (WP_027705705.1), *Pantoea ananatis* (PA13 AER33559.1), *Aspergillus niger* (CAA74744.1), *Zea mays* (ACF85710.1), *Solanum lycopersicum* (AAB09576.1) and *Prunus persica* (AEI70578.1).

Previous cloning and characterisation of *PehA* from *P. carotovorum* reports showed similar results to those previously presented by Hinton et al [37]. In our previous report, the nucleotide sequence of the open reading frame (ORF) of *PehA* gene isolated from *P. carotovorum* showed the sequence identity of 61.5% with *Erwinia tasmaniensis* (WP_042958805.1), 57.8% with *Zymobacter palmae* (WP_027705705.1), and 50.00% with *Pantoea ananatis* (PA13 AER33559.1) respectively [26] which are very close to our present results.

Phylogenetic analysis of the deduced polypeptides encoded by the isolated *PehA* genes compared to other known EndoPGase genes from other microbes, a fungus and of higher plants revealed that *PehA* of *P. carotovorum* individually positioned close to the corresponding EndoPGase of *Pectobacterium brasiliense*. Both *P. carotovorum* and *Pectobacterium brasiliense*. closely positioned with *Erwinia tasmaniensis* since all these bacteria belong to same genus. Similarly, in case of higher plants, *Solanum lycopersicum* and *Prunus persica* positioned close to each other despite of belonging to different families and mutually linked closer to *Zea mays*, a cereal crop. Surprisingly, *Aspergillus niger* was neither closely related to bacterial species nor the higher plants. Similar results were previously presented by Yadav et al [38] who aligned some 48 full-length protein sequences of pectin lyases from varous organisms and generated phylogenetic trees. They found pectin lyases from bacterial and fungal species bifurcating into two distinct clusters. It suggests that bacterial *PehA* are more desirable for transformation of higher plants. The optimal tree with the sum of branch length = 3.71811606 is shown in (Fig 4). The tree is drawn to scale, with branch lengths in the same units as those of the evolutionary distances used to infer the phylogenetic tree.

**Fig 4.** Evolutionary relationships of taxa across microbial and higher plant species for EnoPGase. The values show the evolutionary genetic distances among the species.

3D structure of Isolated EndoPGase was analysed by Phyre2 and is shown in Fig 5.

**Fig 5.** 3D structure of the EndoPGase encoded by the corresponding *PehA* gene expressed in *B. subtilis*. A single-stranded right-handed beta-helix view analysed by Phyre2 and the esidues with active site of the enzyme are highlighted red.

For production of recombinant proteins *Bacillus subtilis* is an attractive host as it is a well-known non-pathogenic organism. *Bacillus subtilis* is considered as GRAS and also ahuge information related to genetic manipulation, protein expression mechanism and fermentation on large scale has been developed for this organism and it will be safer to use in food industries. Hemilä et al [39] expressed the above mentioned *PehA* gene in *Bacillus subtilis* by using a secretion vector with signal sequence and promoter of the gene (*amyE*) encoding α-amylase from *Bacillus amyloliquefaciens*.

For endo-PG expression *Bacillus subtilis* transformants were cultured using 2X-YT medium. Growth Curve of *Bacillus subtilis* in 2X-YT medium is shown in Fig 6. Optical density was measured at 600 nm using microplate. Three sets of competent cells were prepared at stationary phase after every 15 minutes of interval.

**Fig 6.** Growth Curve of *Bacillus subtilis* in 2X-YT medium. Optical density was measured at 600 nm using microplate. Three sets of competent cells were prepared at stationary phase after every 15 minutes of interval.

Recombinant enzyme expression was induced by adding 0.5 mM of IPTG after 2 h and 8 h of inoculation. The enzyme activity was checked periodically after 0, 8, 24, 50 and 72 h of incubation at 37 °C. The maximum activity was observed after 72 h incubation (Fig 7). The culture supernatant was also checked for acitivity by solid plate assay using PG acid and citrus pectin as substrate.

**Fig 7.** Expression recombinant endo-PGase in 2X-YT culture of *Bacillus subtilis*. Medium was initially induced with 10 µl of 0.5 mM IPTG when culture OD reached at 0.8 after 2hrs of inoculation and again induction was done after 8hrs of first induction. Activity of both control (vector clone) and clone (vector with PG gene) was checked after at different time intervals using DNSA method.

The appearance of pink halos with transparent background on polygalacturonic acid plate and clear holes on citrus pectin plates confirmed the activity of recombinant enzyme as shown in Fig 8. The non-concentrated and concentrated recombinant proteins were subjected to SDS-PAGE analysis for molecular identification.

**Fig 8.** Enzyme activity by plate assay. For preparing plate a mixture of 0.8 % agarose and 0.5 % of citrus pectin in sodium acetate buffer of pH 5.0 and dissolved in microwave. Positive control (*Pichia* expression) and *Bacillus* enzyme were filled in the wells, incubated overnight at 37 °C and stained with 0.02 % ruthenium red.

It showed the estimated molecular weight of 48 kDa (Fig 9) corresponding to themolecular mass of endo-PG gene encoding wt-PehA (47 kDa) shown by Massa et al [22]. Previously expression of polygalacturonase encoding *PehA* gene from *Pectobacterium carotovorum* in *E. coli* was reported by Ibrahim *et al* [40]. They reportedthe molecular mass of 41.5 kDa. The pgaA gene encoding endopolygalacturonase expression in *Pichia pastoris* was presented by Liu *et al* [24]. This recombinant enzymeexhibited a molecular weight of 40 kDa and the enzyme assay revealed the optimal pH and temperature of 5.0 and 50 °C respectively.

**Fig 9.** SDS-PAGE analysis of recombinant endo-PGase. Lane 1. Protein marker. Lane 2. Enzyme sample without concentrating. Lane 3. endo-PGase concentrated using Amicon ultra (10K NMWL).

An endo-polyglacturonase gene encoding endo-pga from *Aspergillus niger* was cloned and expressed in *Saccharomyces cerevisiae* EBY100 [41].The report showed molecular mass about 38.8 kDa and optimal conditions for enzyme activity were 50°Cat pH 5.0. Several other reports on cloning and expression of endo-PG into *Pichia pastoris*, other yeasts, and fungi have been published previously. Sawada et al [42] described the cloning of a gene *pehK* encoding exo-Pgase and its expression in *Bacillus subtilis*. Here for the first time we reported cloning and expression of a gene encoding endo-Pgase in *Bacillus subtilis* from *Pectobacterium carotovorum*.

### Biochemical Characterization of Recombinant Enzyme activity

The recombinant enzyme was characterized for optimal pH (Fig 10 (a) and the results showed maximal enzyme activity at pH 5.0. and displayed stability over an extensive pH range from 5.0 to 9.0 (Fig 10 (b).

**Fig 10.** Effect of pH for maximum enzyme activity. **(a)** Enzyme samples were incubated at 40°C for 1 hr in 0.5 % PGA as substrate in different pH buffers. **(b)** .Effect of pH on enzyme stability. Enzyme stability was measured by pre-incubation of sample aliquots in various pH buffers at 25°C for 4 h, then pH was readjusted t to 5.0 and assayed for present activity.

Optimum enzyme activity was observed at a temperature of 40 °C. Fig 11 (a). However, the recombinant enzyme showed an overall decline in stability within the temperature range from 20 °C to 70 °C Fig 11 (b). The recombinant endo-PG retained 80 % of its activity at 30 °C while more than 70 % activity was restored at 40 °C. Zhou *et al* [41] characterized an endo-PgaA from *Aspergillus niger* and expressed in *Saccharomyces cerevisiae*. The recombinant PGase was found active at pH 5.0 and showed a pH stability range within pH 3.0-6.0. The enzyme was active at 50 °C and retained its activity for 60 minute incubation but no activity was observed after 2 h of incubation at 70 °C.

**Fig 11.** Effect of temperature on enzyme activity. **(a)**The substrate and enzyme mixture was incubated at different temperatures for 1 h in 1 mM Sodium acetate buffer(pH 5.0). **(b)**. Effect of temperature on optimum enzyme stability. The enzyme was incubated at various temperatures for 25 min before starting enzyme assay and then the residual activity was measured by conducting standard enzyme assay.

### Effect of Metal Ions

The recombinant enzyme was incubated with 1 mM concentration of metal ions such as Ca^++^, Mg^++^, Zn^++^, EDTA, K^++^ for 45 minutes and residual activity was measured using the standard assay protocol. The graphical figure showed that potassium chloride increased the enzyme activity while EDTA, Zn^++^, Ca^++^, strongly inhibiting the activity (Fig 12). Previously, Li *et al* [43] reported that 1 mM EDTA had no effect on the activity while the same concentration of Ca^++^ was observed to decrease its relative activity.

**Fig 12.** Effect of metal ions on enzyme activity. I mM concentration of metal ions was used in pre-incubation of enzyme-metal mixture at 4 °C for 1 h and residual enzyme activity was measured under standard assay conditions at 40 °C.

An-other report showed inhibition in enzyme activity with the addition of Mg, Zn and Ca [44]. Similarly the activity of an acidic endo-polygalacturonase from *Bispora* sp. MEY-1 [45] expressed in *Pichia pastoris* was strongly inhibited by the addition of EDTA, Zn^++^, Ca^++^, SDS and the activity was not affected by K^++^ and Mg^++^.

## HPLC and TLC analysis

The hydrolyzate of PGA and pectin was analyzed by HPLC and TLC. The reults of TLC revealed that tri and tetra-galacturonates were the end products of substrate hydrolysis (Fig 13 (a). This evidenced the endo-acting action of the recombinant enzyme. While the complex/higher oligosaccharides form a smear on TLC plate and were not further degraded into smaller oligosaccharides. These results confirmed our previously reported analysis of recombinant enzyme from *Pichia pastoris* [26]. HPLC chromatograms (Fig 13(b). obtained from the products of hydrolysis by recombinant EndoPGase showed peaks confirmed the presence of tri and tertra-galacturonates. In comparison with our results recombinant endopolygalacturonase from *Pichia pastoris* and *Saccharomyces cerevisiae* yielded mono, di, and trigalacturonic acids, as the end products of hydrolysis [24, 41].

**Fig 13.** TLC analysis of oigogalacturonates produced by the enzymatic hydrolysis of PGA substrates. **(a)** Lane.1.TLC standard, Lane. 2. Enzyme + 0.5 % PGA, Lane. 3. Enzyme, Lane. 4. PGA 0.5 % + Na-acetate buffer, Lane 5, EndoPGAse + PGA Lane 6, Control. Lane 2 and Lane 5 show the tri and tertra-galacturonates released as a result of enzymatic hydrolysis. **(b)**. HPLC chromatogram showing peaks of tri and tertra-galacturonates as the end products of hydrolyses by the recombinant EndoPGase.

## Conclusions

a. An endo-PGase gene from *Pectobacterium carotovorum* was successfully cloned and expressed in *Bacillus subtilis*. The recombinant enzyme was characterized and release of tri and tetra-galacturonates was observed.
b. The enzyme can be produced on a very simple medium with IPTG induction only.
c. The mode of action of recombinant enzyme was almost similar to other endo-PGase from yeast and fungi. As compared with yeast and fungal expression systems *Bacillus subtilis* might be safe for commercial enzyme preparations.
d. Overall, current study will help in future research works focused to further optimizing the catalytic performance of endopolyglacturonse for processing of pectin-rich materials used in food and fibre industry.

## Competing Interests

**The authors have declared that no competing interests exist**

## Notes

### Competing Interest Statement

The authors have declared no competing interest.

